# Patterns of Cerebellar-Cortical Structural Covariance Mirror Anatomical Connectivity of Sensorimotor and Cognitive Networks

**DOI:** 10.1101/2023.09.28.559942

**Authors:** Zaki Alasmar, M. Mallar Chakravarty, Virginia Penhune, Christopher J. Steele

## Abstract

The cortex and cerebellum are densely connected through reciprocal input/output projections that form segregated loop circuits. Anatomical studies in primates and imaging studies in humans show that anterior lobules of the cerebellum exhibit denser connections to sensorimotor and parietal cortical regions, while lobules Crus I and II are more heavily connected to prefrontal regions. The cerebellum and cortex develop in tandem in childhood, leading to the hypothesis that individual differences in structure should be related, especially for connected regions. To test this hypothesis we examined covariation between the volumes of anterior sensorimotor and lateral cognitive lobules of the cerebellum and measures of cortical thickness (CT) and surface surface area (SA) across the whole brain in a sample of 270 young, healthy adults drawn from the HCP dataset. We found that patterns of cerebellar-cortical covariance differed between sensorimotor and cognitive networks such that anterior motor lobules of the cerebellum showed greater covariance with sensorimotor regions of the cortex, while lobules Crus I and Crus II showed greater covariance with frontal and temporal regions. Interestingly, cerebellar volume showed predominantly negative relationships with CT and predominantly positive relationships with SA. Individual differences in SA are thought to be largely under genetic control while CT is thought to be more malleable by experience. This suggests that cerebellar-cortical covariation for SA may be a more stable feature, whereas covariation for CT may be more affected by development. Additionally, dissimilarity metrics revealed that the pattern of covariance showed a gradual transition between sensorimotor and cognitive lobules, consistent with evidence of functional gradients within the cerebellum. Taken together, these findings are consistent with known patterns of structural and functional connectivity between the cerebellum and cortex. They also shed new light on possibly differing relationships between cerebellar volume and cortical thickness and surface area. Finally, our findings are consistent with the interactive specialization framework which proposes that structurally and functionally connected brain regions develop in concert.

## Introduction

Axonal tracing experiments in primates and neuroimaging studies in humans have revealed an intricate pattern of connectivity between the cerebellum and the cerebral cortex. These two brain structures are connected through reciprocal input/output projections that form loop circuits linking functionally related regions (Leiner et al., 1987; Orioli & Strick, 1989). For example, anterior lobules of the cerebellum exhibit denser connections to sensorimotor and parietal cortical regions than they do to prefrontal regions, while lobules Crus I and II are more heavily connected to the prefrontal regions (Palesi et al., 2017; Salmi et al., 2010). Functional magnetic resonance imaging (fMRI) studies have also revealed functional connectivity between the cortex and cerebellum that parallels structural connectivity, both during rest and during cognitive and motor tasks (Buckner et al., 2011; Kipping et al., 2013). Further, evidence from brain structural development has shown that the cerebellum and cortex develop in tandem in childhood (Kipping et al., 2018). These cerebellar-cortical loop circuits form powerful networks that are thought to be involved in error-processing and the development of forward models that optimize both movement and cognitive functions, such as working memory and language (Guell, Gabrieli, et al., 2018; Schmahmann, 2019).

The pattern of correlations between structural measures can be compared across regions using an approach termed structural covariance (Alexander-Bloch et al., 2013; Lerch et al., 2006). Structural covariance has been examined between structurally connected cortical regions (Mechelli et al., 2005), and between the cortex and subcortical structures, specifically the hippocampus and amygdala (Colibazzi et al., 2007). In addition, recent work from our laboratory has found that the structure of cerebellar and cortical motor regions shows different patterns of covariance in people who began musical training before age seven compared to those who began later and non-musician controls (Shenker et al., 2022). We have hypothesized that this may be a form of interactive specialization where specific experiences, such as music training, promote changes in one region that may drive changes in connected regions (Johnson, 2011). However, and as far as we can determine, normative structural covariance between the cortex and the cerebellum has not yet been examined. Therefore, the goal of the current study was to investigate this relationship in a large sample of healthy young adults from The Human Connectome Project (HCP; Van Essen et al., 2013). Using structural magnetic resonance imaging scans (MRI) of the human brain, we examined the association between the thickness and surface area of the cortex, and the volume of cerebellar lobules. We hypothesized that anatomically connected regions of the cerebellum and cortex would show stronger structural covariance compared to unconnected regions. In particular, we expected that structural covariance would be stronger between posterior frontal and parietal regions with motor lobules of the cerebellum, namely hemispherical lobules III, IV, V, VI, VIIB, VIIIA, VIIIB. We also expected that covariance between prefrontal cortical regions is stronger with lobules Crus I and Crus II, which have been implicated in cognitive functioning and are anatomically connected to frontal regions (Kelly & Strick, 2003).

Anatomical studies in non-human primates have identified differential connectivity between sensorimotor and frontal cortical regions and the cerebellum using transneuronal tracing of fibre pathways (Kelly & Strick, 2003). Axons from lobules III-VI as well as lobules VIIIA, and VIIIB were found to project to the primary motor cortex, the premotor cortex, and supplementary motor cortex. Importantly, another set of axons projects back from these cortices to the cerebellum to form cerebellar-cortical loops. Similarly, Crus I and Crus II have been found to be connected to frontal regions, primarily the dorsal prefrontal cortex, through reciprocal loops.

As expected, cerebellar-cortical functional connectivity reflects the known structural connectivity, as observed in resting-state fMRI (rs-fMRI) and latent functional gradients (Guell, Schmahmann, et al., 2018; Stoodley & Schmahmann, 2010). Observed patterns of rs-fMRI connectivity between the cerebellum and sensorimotor and prefrontal regions match the anatomical connections between the regions described above (Buckner et al., 2011). Resting-state connectivity, which measures correlated activity between brain regions, has been found between the lobules of the anterior cerebellum, VIIIA, and VIIIB and sensorimotor cortices, and between Crus I and Crus II and the prefrontal cortex (Kipping et al., 2013). Consistent with the results of structural and functional connectivity studies, task-based studies show that lobules III-VI, VIIA-VIIIB, and sensorimotor cortices are active during sensorimotor tasks and Crus I and Crus II and the prefrontal cortex are active during cognitive functions (Buckner et al., 2011; Salmi et al., 2010). At this functional level, connections between the cerebellum and cortex have been proposed to underlie cerebellar contributions to optimizing motor and cognitive behaviour (Balsters & Ramnani, 2008; Ito, 2005; Roth et al., 2013). Indeed, correlated activity between sensorimotor cortices and the cerebellum has been shown to be related to motor task performance and motor learning (Penhune & Steele, 2012; Stoodley & Schmahmann, 2010), and correlated cerebellar-cortical activity has also been linked to executive functioning (Stoodley et al., 2012). Thus, overlapping functional and structural connectivity between the cortex and the cerebellum supports the idea that the two regions interact to optimize performance in a wide range of cognitive and motor functions. Gradient decomposition of cerebellar function also supports this conclusion by revealing motor vs non-motor representations within the cerebellum (Guell, Schmahmann, et al., 2018). These representations are consistent with the cortical gradient organization from primary to transmodel regions (Margulies et al., 2016). However, whether the associations between the structure of the cerebellum and that of the cortex show similar organization remains unclear.

We and others hypothesize that the structural and functional connectivity between the cortex and the cerebellum may be in part determined by a pattern of interactive changes during development (Johnson, 2011). Johnson (2011) proposes that functionally connected brain regions develop in association and exert effects on each other throughout maturation and in response to experience. This framework for understanding brain development is termed interactive specialization. Based on the existence of structural and functional connections between regions (Alexander-Bloch et al., 2013), interactive specialization might be expected to lead to structural covariance. Structural connectivity has been suggested to mediate positive associations between variations in the structure of brain regions during development (Mechelli et al., 2005). Structural covariance between regions is also influenced by genetics and lifespan experiences and exhibits different patterns across sexes (Chen et al., 2011; Lv et al., 2010; Schmitt et al., 2008). It has been shown that dancers exhibit an association between reduced cortical thickness in the entire brain and cortical thickness in middle frontal gyrus when compared to healthy controls(Karpati et al., 2018). Yee et al. (2018) observed an association between the transcriptomic similarity between regions and their structural covariance, reflecting a genetic influence on these patterns of variation. Since structural covariance is influenced by life experiences, we expect that experience could play a role in cerebellar-cortical structural associations as well. This is evident in our previous study with musicians showing that changes in cerebellar-cortical structural covariance were linked to musical training before the age of seven (Shenker et al., 2022). The current study extends this work to investigate the patterns of normative cerebellar-cortical structural covariance in the healthy adult brain.

There are two primary and theoretically distinct measures of cortical structure that can be measured in humans from MRI: cortical thickness and surface area. Cortical thickness is defined as the depth/thickness of the grey matter ribbon, and surface area as the 2-dimensional extent of a given region of cortex. Individual variation in these features is thought to reflect different contributions from genetic and environmental factors during development (Panizzon et al., 2009). Surface area undergoes rapid changes early in life and is thought to be under greater genetic control (Bishop et al., 2000; Sanabria-Diaz et al., 2010; Yoon et al., 2012). Cortical thickness continues to change across development into young adulthood and is therefore more likely to be affected by the environment and experience (Amlien et al., 2016). While cortical thickness and surface area are routinely extracted from MRI images of cortex, measuring them in the cerebellum is problematic. The cerebellum has a relatively thin grey matter ribbon, very dense gyrification, and closely packed lobules. This means that white and grey matter tissue segmentation, the basis for estimating these metrics, is unreliable (Sereno et al., 2020). Therefore, lobular volume is a more reliable measure of cortical structure in the cerebellum that can be accurately measured. To measure the volumes of cerebellar lobules, we used a robust multi-atlas segmentation approach (Chakravarty et al., 2013) that has been applied in a number of previous studies of cerebellar structural variation (Park et al., 2014; Shenker et al., 2022; Steele & Chakravarty, 2018).

Taken together, the existing literature suggests that there may be normative covariance between the structure of the cortex and cerebellum based on known anatomical and functional connectivity. Therefore, the current study examined cerebellar-cortical structural covariance in a large sample of healthy young adults controlling for age and sex. We hypothesized that the volumes of motor and cognitive lobules of the cerebellum would show greater covariance with cortical thickness and surface area of motor and cognitive regions of the cortex. Further, we expected that these patterns of covariation might differ for cortical thickness and surface area, and between hemispheres.

## Method

### Participants

Structural MRI images were obtained from the Human Connectome Project S1200 release (HCP; full protocol is described in (Van Essen et al., 2013)). We conducted our analyses on a subsample of 270 right-handed young adults (age: *M:* 28.77 years, *SD:* 3.73; 157 female) with no history of psychiatric, neurological, or neuropsychological disorders, and no history of substance abuse. A previous study reported the volumes of all cerebellar regions in this same sample (Steele & Chakravarty, 2018).

### Procedure

#### MRI Acquisition and Preprocessing

T1-weighted Structural MRI images were acquired for all participants at the WashU research facilities on a 3T Connectome Skyra MRI scanner with a 32-channel head coil. Scanning parameters were as follows: voxel size = 0.7 mm isotropic, reptation time = 2400 ms, inversion time = 1000 ms, echo time = 2.14 ms, field of view = 224×224 mm (Van Essen et al., 2012). All MRI scans were preprocessed by the HCP according to the minimal preprocessing pipeline (Glasser et al., 2013). This included registration to the common Montreal Neurological Institute 152 space (MNI-152) with a rigid body transformation. Then, FreeSurfer’s recon-all was used to reconstruct white and grey matter surfaces that were then used in surface area (SA) and cortical thickness (CT) measurements (Dale et al., 1999).

#### Thickness and Area calculation

Thickness was measured as the distance between the white and gray matter surfaces in millimeters (mm). To extract the SA of the cortex at every vertex, the area of one third of each triangle that the vertex was part of was assigned to that vertex, and it was measured in mm^2^. Surface area values for each participant were normalized at each vertex by total brain volume to account for global differences in brain size. SA, but not CT values were normalized because variations in the cortical volume are mainly driven by changes in SA, therefore normalizing only SA corrects for brain size while preventing data contamination with noise (Im et al., 2008).

Thickness and area measurements were then smoothed using a 12mm full width at half maximum smoothing gaussian kernel.

#### Cerebellar Lobule Identification

To label the cerebellar lobules of interest, the Multiple Automatically Generated Templates Brain Segmentation Algorithm was used (MAGeTbrain: https://github.com/CobraLab/MAGeTbrain; (Park et al., 2014)). The algorithm first performs a non-linear registration between multiple high-resolution, expert-labelled, atlases and a set of 21 randomly chosen scans from the subsample (Steele & Chakravarty, 2018). Each of the 21 scans are segmented according to the 5 atlases, therefore each voxel in these scans would have 5 labels assigned to it. Then, a majority vote is conducted to determine the most frequent label out of the 5 × 21 = 105 possible labels that each voxel is assigned. Based on the votes, a sample template is created, and each participant’s scan was then labelled according to that template. Following the identification of the nine hemispherical cerebellar lobules in each hemisphere (III, IV, V, VI, Crus I, Crus I, VIIB, VIIIA, VIIIB), we normalized their volume by total brain volume to account for variation due to brain size.

### Statistical Analysis

#### Cerebellar-cortical Structural covariance

Since our goal was to examine cerebellar-cortical structural covariance, regression analyses were used to investigate the relationships between measures of CT and SA at each vertex in the cortex and the volumes of contralateral cerebellar lobules. For these analyses, cerebellar lobular volumes were the independent variables and either CT or SA served as the dependent variables. Age and sex were included as covariates. Standardized regression coefficients were extracted to indicate the specific association between the volumes of the lobules and vertex-wise CT or SA, which were then were plotted on the cortex for visualization. In total, we conducted 18 regression analyses at each of the 326k cortical vertices (i.e., 9 lobules each for CT and SA, with each lobular volume serving as the independent variable).

#### Dissimilarity Assessment

To assess dissimilarity in patterns of associations between the cortex and the volumes of the different cerebellar lobules, we computed the Euclidean distance between the vertex-wise associations of each cortical hemisphere (for CT and SA separately) for each cerebellar lobule. In this step, Euclidean distance represents the dissimilarity in the pattern of covariance between each pair of lobules, and provides a comparative summary of the vertex-wise regression analyses. In this analysis, greater distance between lobules is indicative of greater dissimilarity, whereas lower values indicate similar patterns of covariance of the lobules with CT/SA across the cortical hemisphere. We conducted 36 dissimilarity assessments (each of the 9 lobules to every other lobule) per cortical measure per hemisphere. In total, there were 36 × 4 = 144 assessments, grouped by measure (CT/SA) and hemisphere.

## Results

### Cerebellar-cortical Structural covariance

First, we assessed cerebellar-cortical structural covariance between motor and cognitive lobules of the cerebellum and either CT or SA using multiple regression accounting for age and sex. Results revealed a pattern of cerebellar-cortical covariance that differed for CT and SA, between sensorimotor vs cognitive regions, and between the left and right hemispheres (Figure. 1).

**Fig.1.**
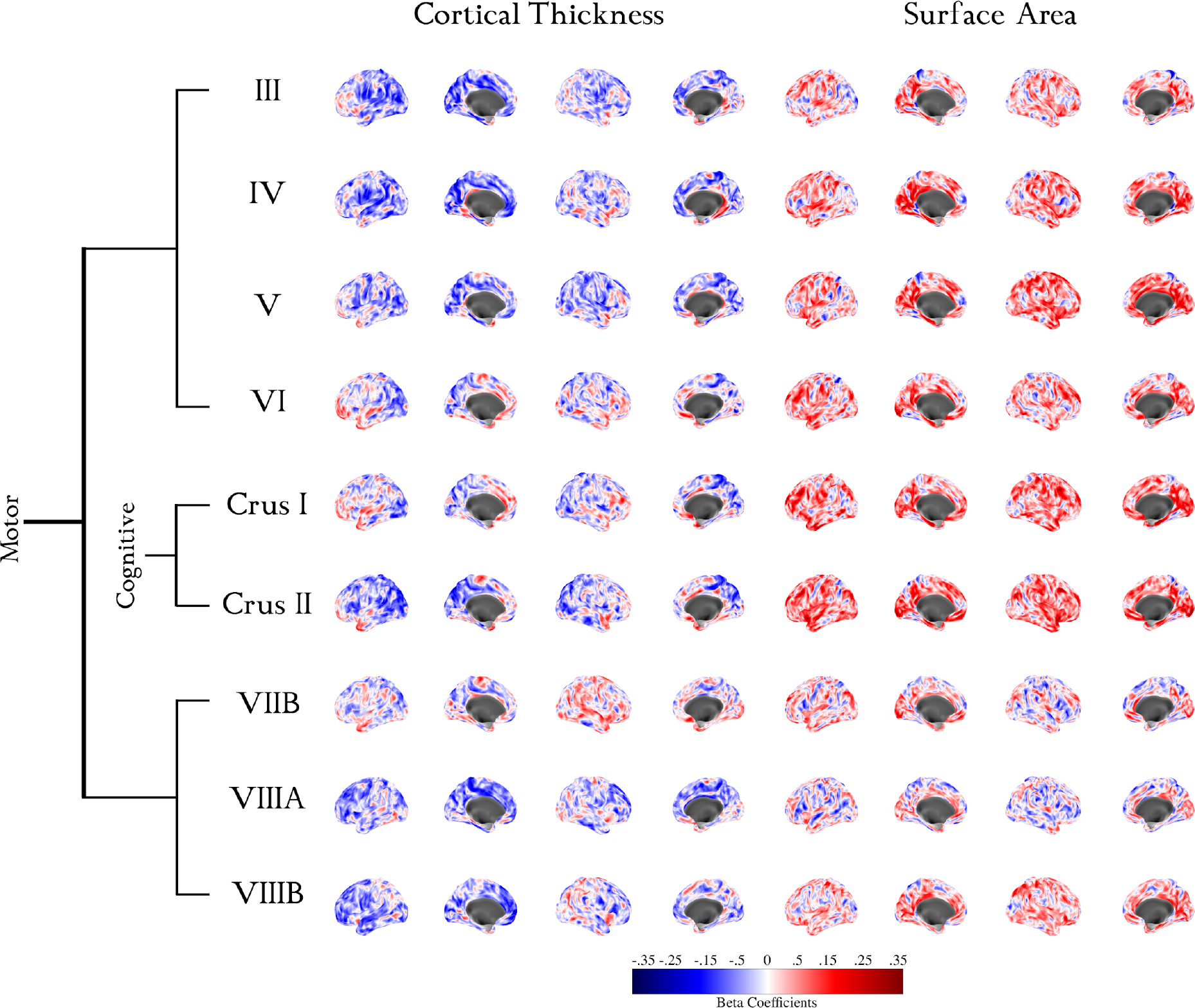
vertex-wise beta coefficients of the association between CT and SA of the cortex and the volumes of cerebellar lobules while accounting for age and sex. Each brain-plot represents the association between the cortical measure (top) of one hemisphere (lateral and medial view) and the volume of the contralateral cerebellar lobule (right). Blue indicates a negative association while red indicates a positive one.

For CT, covariance between left cortical sensorimotor regions and right lobules III, IV, V, VIIIA, and VIIIB were predominantly strong negative associations as seen by the large negative beta coefficients. This negative covariance was weaker for lobules Crus I and Crus II, and positive for lobules that are at the boundary of motor and cognitive transitions, namely VI and VIIB. CT of the left cortical cognitive regions in the frontal lobes exhibited weaker negative associations with the motor lobules III, IV, V, VIIIA, and VIIIB, and strong positive associations with lobules VI, Crus I, Crus II, and VIIB. Overall, this pattern was weaker and less consistent for the right hemisphere. For CT of the right cortical sensorimotor regions, negative associations were found with lobules III, IV, V, while lobules VI – VIIIB show weak positive associations. CT of right frontal regions was positively associated with lobules VI – VIIB. In contrast to the left cortex, CT of the right frontal regions was also weakly positively associated with lobules III – VIIB, and negatively associated with VIIIA and VIIIB.

In contrast to CT, SA was generally positively related to cerebellar volumes, but with a similar pattern in the shift of covariance across cognitive vs motor lobules and hemisphere. SA of the left sensorimotor regions of the cortex were strongly positively associated with right lobules III, IV, and V. This positive association extended to lobules VI, Crus I, Crus II, unlike CT where the direction of association switched from negative to positive between motor and cognitive lobules. We observed negative associations between left SA and right lobules VIIB and VIIIA in a similar fashion to lobules III and IV. However, the associations between right VIIIB and SA were predominantly positive. SA of left frontal regions was strongly positively associated with lobules Crus I and Crus II, with weaker associations with other lobules. A similar pattern of associations was observed for the right hemisphere, where SA was positively associated with volumes of lobules III – Crus II as well as VIIIB, and negatively associated with VIIB and VIIIA.

### Dissimilarity

To quantify differences in cerebellar-cortical structural covariance between motor and cognitive regions, we investigated whether the distributions of covariance were different across lobules for CT and SA. As a measure of dissimilarity, we calculated the Euclidean distance between hemispherical covariance maps for each pair of lobules (Figure 2). Each association map is compared to other maps in the same hemisphere for either CT or SA, summarizing vertex-wise dissimilarity between each pair of lobules in one score.

**Fig.2.**
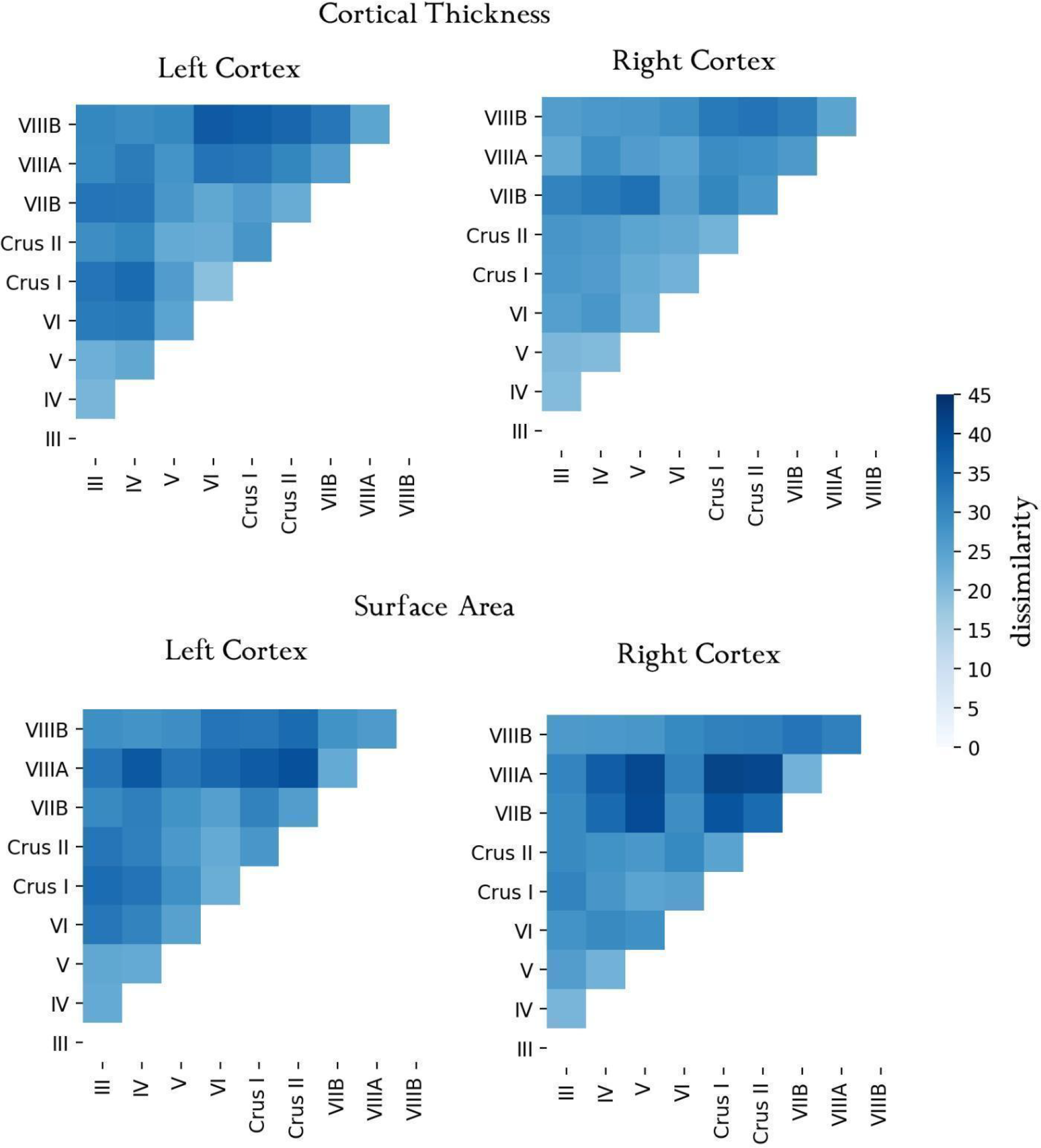
Euclidean distance between each pair of cerebellar-cortical associations, for each structural measure of each hemisphere. The distance represents the dissimilarity between each brain-plots in figure 1 and all the other plots below it, for all possible pairs.

We observed a clear delineation of motor vs cognitive cerebellar lobules in their associations with the cortex (Figure 2). For CT, motor lobules III-V showed higher similarity among themselves and greater dissimilarity with lobules VI, Crus I and VIIB. Cognitive lobules Crus I and II also showed higher similarity with each other and greater dissimilarity with lobules III-V and VIIIA and VIIIB. Lobule VI appears to be a transitional area, showing less dissimilarity with Crus I and II (than the other motor lobules) and greater dissimilarity with other motor lobules. Dissimilarity was generally greater in the left than the right hemisphere.

The dissimilarity between cognitive and motor lobules was overall greater for SA. Motor lobules III-V again showed higher similarity with each other and greater dissimilarity with lobules Crus I and II, and VIIIA. Cognitive lobules Crus I and II were more similar with each other and showed greater dissimilarity with lobules VIIB and VIIIA. Lobule VI again was found to be more similar to Crus I and II and more dissimilar to lobules VIIB and VIIIA. Inferior motor lobules VIIB and VIIIA also showed marked dissimilarity from all other regions. For SA, dissimilarity appears to be more pronounced for the right, compared to the left hemisphere.

## Discussion

The goal of the current study was to examine the pattern of cerebellar-cortical structural covariance in motor and cognitive networks. Overall, our findings revealed a pattern of cerebellar-cortical covariance that differed between sensorimotor vs cognitive regions, for cortical thickness and surface area, and between the left and right hemispheres. Cortical thickness showed predominantly negative relationships with cerebellar volume, particularly in sensorimotor, parietal and frontal regions of the left hemisphere. Surface area showed a predominantly positive relationship with cerebellar volumes, particularly in sensorimotor, parietal, frontal and temporal lobe regions bilaterally. Motor lobules (III-VI & VIIB-VIIIB) of the cerebellum showed greater covariance with sensorimotor regions of the cortex, and cognitive lobules (Crus I & Crus II) showed greater covariance with frontal and temporal regions. The pattern of cerebellar-cortical covariance for both cortical thickness and surface area differed across regions, with covariance being more similar within sensorimotor and cognitive networks and more dissimilar between them. Dissimilarity in the pattern of covariance of each lobule with the cortex showed a gradual transition between sensorimotor and cognitive lobules, with lobules III-VI showing the greatest dissimilarity with lobules VI and VIIB showing greater similarity to Crus I and II. Taken together, these findings are consistent with the known pattern of anatomical connectivity between different cerebellar lobules and the cortex. In addition, they shed new light on the relationship between cerebellar volumes and different cortical features, showing that larger cerebellar volumes are related to reduced cortical thickness but greater surface area. Surface area has been hypothesized to be under greater genetic control (Bishop et al., 2000; Sanabria-Diaz et al., 2010; Yoon et al., 2012) while cortical thickness is thought to be more malleable by experience (Amlien et al., 2016). This suggests that covariance between cerebellar volumes and surface may be a more stable feature, whereas covariance with cortical thickness may be more affected by development. This is consistent with the interactive specialization framework which proposes that functionally connected brain regions develop in tandem (Johnson, 2011).

For both the left and right hemisphere, cerebellar motor lobules III-V and VIIIA and B showed primarily negative associations with sensorimotor and parietal cortical regions. In contrast, lobules VI and Crus I showed positive associations with frontal regions. In a parallel finding, but with reversed direction, motor lobules III – V and VIIB, VIIIA and B showed predominantly positive associations with cortical surface area in sensorimotor regions of the cortex. The volumes of Crus I and II were strongly positively associated with surface area of cortical frontal regions. The pattern of observed differences in structural covariance were supported by a measure of dissimilarity (Euclidean distance) which shows clearly that cerebellar-cortical covariance within motor and cognitive regions is more similar than covariance across these regions, and that dissimilarity increased at the transition from motor (III and IV) to cognitive (Crus I and Crus II) and back to motor (VIIIA and VIIIB) lobules.

In line with our hypotheses, these differing patterns of cerebellar-cortical structural covariance are consistent with known anatomical connectivity in primates (Kelly & Strick, 2003), with studies in humans using diffusion-weighted imaging (Habas & Cabanis, 2007; Rousseau et al., 2022; Steele & Chakravarty, 2018) and with evidence from resting-state functional connectivity (Bernard et al., 2012; Krienen & Buckner, 2009; Stoodley & Schmahmann, 2010). In primates, lobules III-VI as well as VIIIA and VIIIB project primarily to motor cortical regions, while Crus I and II connect primarily to prefrontal regions (Kelly & Strick, 2003). Evidence for a similar pattern of connectivity has been found using diffusion imaging in humans, showing distinct motor and non-motor divisions in the dentate nucleus and the pons (Rousseau et al., 2022; Steele et al., 2017). In parallel, patterns of resting-state functional connectivity between the cerebellum and sensorimotor and prefrontal regions match the anatomical connections between the regions described above (Buckner et al., 2011; Ji et al., 2019; Marek et al., 2018; O’Reilly et al., 2010). Finally, task-based studies show specific activation of sensorimotor regions of cerebellum and cortex for sensorimotor tasks and lobules Crus I and Crus II and the prefrontal cortex for cognitive tasks (King et al., 2019, 2023; Salmi et al., 2010; Stoodley et al., 2012).

Thus the observed pattern of structural covariance is consistent with known segregation of the sensorimotor and cognitive cortico-cerebellar networks. This segregation is also supported by the results of the dissimilarity metric, which showed that covariation was more similar within motor and cognitive networks than between them. Despite this broad segregation into motor and cognitive domains, we also observed a transitional zone at the border of motor to cognitive representations for lobules VI and VIIB. This is consistent with the observation of multiple gradients of functionally-defined continuous representations that do not strictly match lobular structural boundaries (Guell & Schmahmann, 2020; King et al., 2019).

An intriguing finding from this study is the differing relationship between cerebellar volumes and the cortical metrics; while cerebellar volumes exhibited largely negative relationships with cortical thickness, they showed largely positive relationships with surface area. These opposite relationships may be due to differing developmental trajectories: surface area increases up until age 12 with little change thereafter, while cortical thickness peaks in infancy and then decreases until the third decade of life (Amlien et al., 2016; Bethlehem et al., 2022). Based on this, variation in surface area is thought to be largely under genetic control, while variation in cortical thickness is thought to reflect a combination of accumulated genetic and environmental effects (Roe et al., 2023). Together with our findings, this suggests that interactions between cerebellar and cortical regions across development may lead to correlated structural plasticity, although it is not clear what underlying mechanisms would lead to positively correlated versus negatively correlated changes.

Structural covariance is thought to be driven by structural and functional connectivity (Alexander-Bloch et al., 2013). Therefore, covariation between cerebellar and cortical regions is consistent with the idea that functionally and structurally connected regions may develop in tandem, a concept known as interactive specialization (Johnson, 2011). Developmental data show that GM volume of anterior motor regions, including M1 and PMC has a peak rate of change between the ages of 6 and 8 (Bethlehem et al., 2022; Giedd et al., 1999). In contrast, peak maturation in the cerebellum occurs later, between the ages of 12 and 18 (Bethlehem et al., 2022; Tiemeier et al., 2010). The interactive specialization framework proposes that connected brain regions or networks interact during development to reciprocally influence maturation (Johnson, 2011). In the case of the cortex and cerebellum, we propose that dense connectivity drives plasticity, and that the structure of these regions changes interactively (Fjell et al., 2019; Penhune, 2020). The cerebellum has been hypothesized to support optimization of both motor and cognitive functions through its loop circuits with different cortical regions (Bostan et al., 2013), possibly through encoding of predictions and error correction (Sokolov et al., 2017). Thus optimal functioning is contingent on the co-development of the cortex and the cerebellum. This idea is supported both by correlated cerebellar and cortical structural changes in early-trained musicians (Shenker et al., 2022) and in rodents exposed to an enriched environment (Scholz, Allemang-Grand, et al., 2015; Scholz, Niibori, et al., 2015). Earlier-maturing sensorimotor cortical networks may modulate development of later-maturing cerebellar networks. Later in development, however, cerebellar mechanisms related to forward models and optimization may contribute to the fine-tuning of motor and cognitive functions observed in adolescence and early adulthood (Fuhrmann et al., 2015).

Finally, our results also show that the degree of covariance was overall stronger for the left compared to the right hemisphere. This may be due to typical left-hemisphere dominance for control of the right hand and language functions. It may also be due to differences in intra- and interhemispheric connectivity where the left hemisphere is characterized by stronger within hemisphere networks and the right by broader, more distributed connections (Iturria-Medina et al., 2011).

Although we aimed to decompose cortical volumes into more basic elements by using cortical thickness and surface area, it remains difficult to attribute the patterns of structural covariance that we observed to specific molecular or physiological causes. Further research is needed to examine the mechanisms that drive structural covariance or determine the direction of correlated change. One approach that could address this would be to look at covariance in a longitudinal sample of children where patterns of change in cerebellar volume could be related to change in cortical metrics across development. It is also likely that the observed patterns of cerebellar-cortical structural covariance would differ with aging since surface area and cortical thickness follow different trajectories (Lemaitre et al., 2012). Another direction for future work would be to decompose cerebellar volumes into surface area and cortical thickness in order to examine covariation with cortex for each metric separately. While we were able to estimate lobular volumes with high accuracy, the cerebellar cortex is a dense and highly gyrified structure, rendering results from commonly used tools to measure cortical thickness and surface unreliable (Sereno et al., 2020). In the future, use of high-field strength scanners may resolve these limitations.

## Conclusion

The observed pattern of cerebellar-cortical structural covariance largely mirrors known structural and functional connectivity, where sensorimotor and cognitive regions are connected in partially separate loops. Dissimilarity metrics also identified transitional regions at the boundaries of motor and cognitive lobules, consistent with evidence of functional gradients, rather than sharp boundaries within the cerebellum. Patterns of structural covariation differed for surface area and cortical thickness, consistent with evidence that individual variation in these features results from either greater genetic or environmental influences. Together, these findings support the interactive specialization framework which proposes that structurally and functionally connected regions mutually influence each other during development.

